# Generation of an Unbiased Interactome for the Tetratricopeptide Repeat Domain of O-GlcNAc Transferase Indicates a Role for the Enzyme in Intellectual Disability

**DOI:** 10.1101/2020.07.30.229930

**Authors:** Hannah M. Stephen, Jeremy L. Praissman, Lance Wells

## Abstract

The O-GlcNAc transferase (OGT) is localized to the nucleus and cytoplasm where it regulates nucleocytoplasmic proteins by modifying serine and threonine residues with a non-extended monosaccharide, β-N-Acetyl-Glucosamine (O-GlcNAc). With thousands of known O-GlcNAc modified proteins but only one OGT encoded in the mammalian genome, a prevailing question is how OGT selects its substrates. Prior work has indicated that the N-terminal tetratricopeptide repeat (TPR) domain of OGT, rather than its C-terminal catalytic domain, is responsible for subcellular targeting and substrate selection. An additional impetus for exploring the OGT TPR domain interactome is the fact that missense mutations in *OGT* associated with X-linked intellectual disability (XLID) are primarily localized to the TPR domain without substantial impact on activity or stability of the enzyme. Therefore, we adapted the BioID labeling method to identify interactors of a TPR-BirA* fusion protein in HeLa cells. We identified 115 high confidence interactors representing both known and novel O-GlcNAc modified proteins and OGT interactors. The TPR interactors are highly enriched in processes in which OGT has a known role (e.g. chromatin remodeling, cellular survival of heat stress, circadian rhythm), as well as processes i n which OGT has yet to be implicated (e.g. pre-mRNA processing). Importantly, the identified TPR interactors are involved in several disease states but most notably are highly enriched in pathologies featuring intellectual disability. These proteins represent candidate interactors that may underlie the mechanism by which mutations in *OGT* lead to XLID. Furthermore, the identified interactors provide additional evidence of the importance of the TPR domain for OGT targeting and/or substrate selection. Thus, this defined interactome for the TPR domain of OGT serves as a jumping off point for future research exploring the role of OGT, the TPR domain, and its protein interactors in multiple cellular processes and disease mechanisms, including intellectual disability.

## Introduction

The O-GlcNAc transferase (OGT) is a nucleocytoplasmic glycosyltransferase that modifies substrate proteins with a β-N-acetylglucosamine (O-GlcNAc) on serine and threonine residues. OGT is a unique mammalian glycosyltransferase in that it modifies intracellular proteins outside of the secretory pathway, and the O-GlcNAc modification it creates is non-extended, dynamic, and inducible^1^. The O-GlcNAc modification is often compared to phosphorylation, given their similar characteristics and the fact that both occur on thousands of nuclear and cytosolic proteins^2^. In fact, OGT and Ser/Thr kinases often compete for the same sites on certain protein substrates^2^ and can regulate each other by post-translational modification^3,4^. However, unlike protein phosphorylation which is mediated by hundreds of kinases, there is only one gene encoding intracellular O-GlcNAc Transferase in mammals. Thus, not surprisingly, OGT is essential for life^5^ and is involved in many intracellular processes including nutrient sensing, transcription, and cellular stress^1^. OGT also has been implicated in many diseases including cancer, Alzheimer’s disease, diabetes, and more recently, in X-Linked Intellectual Disability (XLID) as identified originally by our team in collaboration with clinical partners^6^ and further confirmed and expanded on by our group and others^7–9^.

Given the wide diversity of OGT substrates and functions, and the existence of only one mammalian OGT, a prevailing question in the O-GlcNAc field is how OGT selects its substrates. Previous research suggests that the N-terminal tetratricopeptide repeat (TPR) domain of OGT (consisting of 13.5 repeats in the full-length protein), rather than its C-terminal catalytic domain, is responsible for OGT substrate selectivity^10–12^. However, the hypothesis of the TPR domain mediating protein-protein interactions has only been directly tested for a few select proteins^13–16^. A few early attempts to define the full-length OGT-interactome using co-immunoprecipitation have also been made^17,18^. An unbiased approach to identifying proteins that interact specifically with the TPR domain would lend further support to the hypothesis of the TPR domain mediating OGT substrate selectivity, and allow for the identification of new potential substrates and “partner proteins”, which interact with the TPR domain of OGT to target it to specific subcellular regions and/or protein complexes.

An additional impetus for TPR interaction studies is the observation that the majority of reported missense mutations in *OGT* causal for XLID are localized to the TPR domain and do not grossly affect catalytic activity or stability, suggesting a potential protein-protein interaction-based mechanism^6,7^. Therefore, to demonstrate that the TPR domain of OGT is capable of substrate selection, and to capture endogenous OGT TPR interactors including transient interactors, we took advantage of the BioID method, utilizing a fusion protein consisting of the full-length OGT TPR domain with a modified biotin ligase in place of the catalytic domain of OGT.

BioID is a well-established proximity proteomic labeling method that utilizes a promiscuous biotin ligase (BirA*) to label nearby proteins with biotin, which allows them to easily be extracted and identified using mass-spectrometry based proteomics^19^. Using a TPR-BirA* fusion protein in a HeLa cell system (and a eGFP-BirA* fusion protein as a negative control), we have identified over 100 high-confidence OGT TPR interactors, including both known and novel OGT substrates and interactors. This work strongly suggests that the TPR domain, through protein-protein interactions, plays a major role in OGT substrate selectivity. Exploiting these interactions may allow for fine-tuning of the modification of specific O-GlcNAc modified substrates which has been explored using other techniques^20,21^. This protein set also further confirms OGT’s role in many cellular processes and reveals potential novel pathways in which O-GlcNAc may play an intricate role. Finally, the interactome is highly enriched in proteins involved in neurological disorders that present with intellectual disability. These proteins represent a set of candidate interactors to explore for future mechanistic studies of the functional role of OGT and the O-GlcNAc modification in XLID.

## Experimental Procedures

### Plasmid Constructs

Plasmids for proximity proteomics were constructed on a CMV promoter with a C-terminal BirA*. Fusion gene construction is as follows. For TPR BirA*: TPR – 3X GGGGS linker – BirA* – 2X FLAG. For eGFP-BirA*: eGFP – 3X GGGGS Linker – BirA* – 2XFLAG. The BirA* sequence was obtained from the original paper describing BioID^19^. For TPR-BirA*, residues 1-473 of OGT (consisting of the 13.5 TPR repeats of full-length OGT, Uniprot Accession O15294) were used. Full fusion protein sequences are in Supplementary Table 1.

### Cell culture and BioID

HeLa cells were grown in DMEM with 10% FBS on 14.5cm plates, passaged for maintenance every 4-6 days (1:4-1:10). For expression of fusion proteins, cells were transfected at ~70% confluency using lipofectamine 2000 (Invitrogen) according to manufacturer recommended ratios with 60μg plasmid DNA for TPR-BirA* and 6μg plasmid DNA for eGFP-BirA* (eGFP-BirA* expresses at a much higher level than TPR-BirA* – see **Fig. 1 C/D**). After 24 hours, cell media was replaced with media containing 50uM biotin to induce labeling for 24 hours. Cells were then collected and subjected to nucleocytoplasmic lysis to collect protein. Briefly, cells were lysed in hypotonic buffer A (10mM Tris-HCl pH 5.5, 500uM DTT, 500uM EDTA, protease and phosphatase inhibitor cocktails (Sigma)) for 10 minutes, vortexed twice for 30 seconds, then lysed in hypertonic buffer B (10mM Tris-HCl pH 7.5, 800mM NaCl, protease and phosphatase inhibitors, and universal nuclease (ThermoFisher Scientific)) for 5 minutes, followed by 30 seconds of highspeed vortexing and an additional 5 minute incubation. Membranes were pelleted by centrifugation, and the supernatant containing cytosolic and nuclear proteins used in further analysis.

**Figure 1:**
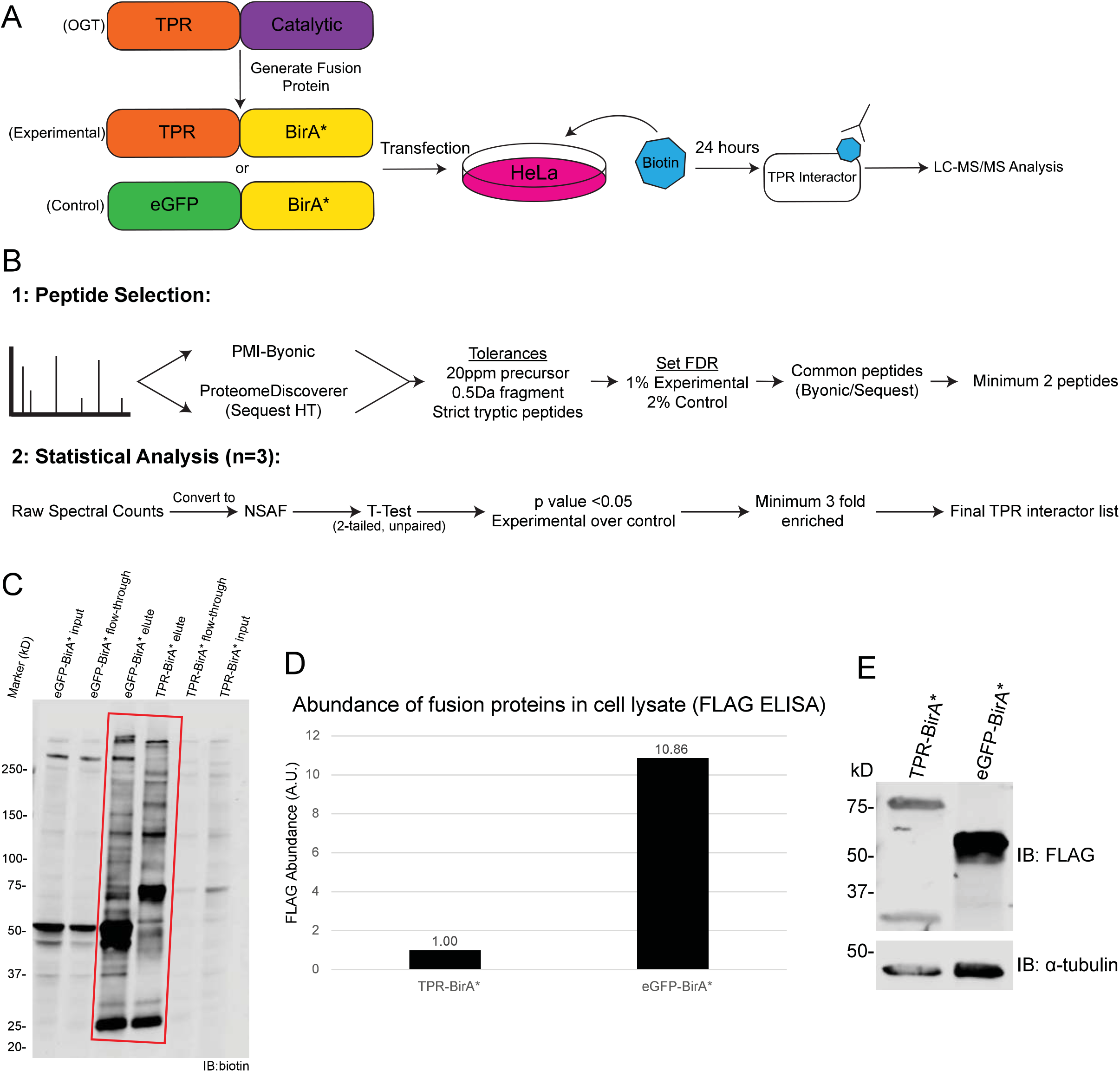
BioID approach to define OGT TPR interactors in HeLa cells. ***A***, Workflow of BioID process for identifying TPR interactors. ***B***, Workflow of MS analysis and protein validation. Samples were prepped in biological triplicate. ***C***, Western blot showing expression of TPR-BirA* and eGFP-BirA* fusion proteins (anti-FLAG tag), α-tubulin as loading control. 20ug/lane. ***D***, ELISA quantification of FLAG-tagged fusion proteins (one representative replicate, A.U.=Arbitrary units) ***E***, Representative western blot with anti-biotin antibody demonstrating enrichment of biotinylated proteins following biotin immunoprecipitation (representative blot from three replicates). Red box indicates lanes showing enrichment of biotinylated proteins by biotin IP. 10μg/lane. FT=flow-through. For elute and FT, 10ug/lane. For elute, 25% of total eluate loaded.

### Quantification of FLAG-tagged proteins

FLAG-tagged fusion proteins were quantified from HeLa cell lysate using a FLAG ELISA (Cayman Chemical) according to the manufacturer’s protocol.

### Identification of biotinylated proteins

Biotinylated proteins were purified from cellular lysate using an anti-biotin immunoprecipitation protocol as previously described^22^. 25% of eluate was reserved for anti-biotin western blot, then remaining biotinylated proteins were then run out on a 10% SDS-PAGE gel using the Bio-Rad Mini-PROTEAN gel system. The gels were not stained. Each lane was cut into four equal fractions based on molecular weight markers, then extracted, reduced, alkylated, and digested with trypsin as previously described^22^. Dried peptides were resuspended in 40μL sample buffer (10% Buffer B (80% acetonitrile, 0.1% formic acid), 90% Buffer A (0.1% formic acid), and 15uL of this was injected for each instrument run. LC-MS/MS was performed on an Orbitrap Fusion Tribrid mass spectrometer (ThermoFisher Scientific) equipped with an Ultimate 3000 RSLCnano HPLC system (Thermofisher Scientific). Peptides were separated on an Acclaim™ PepMap™ RSLC C18 column (75 μm ID × 15 cm; 2 μm particle size) at a flow rate of 0.200μL/min over a 150min linear gradient of 1-99% Buffer B with a total run time of 180min. Precursor scans were collected using the Orbitrap mass analyzer with a scan range of 300-2000m/z and mass resolution of 60,000. Most intense ions were fragmented using 38% CID collision energy and detected in the Ion Trap with 1 microscan and dynamic exclusion for 15 seconds after one occurrence. Samples were run in randomized pairs within a replicate, with each pair consisting of a corresponding gel fraction in eGFP-BirA*, run first, then TPR-BirA*, run second, with one 90 min wash in between each sample and two 90 minute washes in between pairs (20uL 10% Buffer B injection for washes). The raw data for all 24 LC-MS/MS analyses (12 control and 12 experimental) has been deposited to the MassIVE database (https://massive.ucsd.edu/ProteoSAFe/static/massive.jsp, Dataset ID: MSV000085626).

### Experimental design and statistical rationale

Three biological replicates were performed, each consisting of one TPR-BirA* and one eGFP-BirA* sample, with TPR-BirA* representing the experimental condition and eGFP-BirA* representing the negative control. An n of 3 was selected to allow us to perform statistical analyses with sufficient statistical power. Spectral counts were converted to normalized spectral abundance factors (NSAFs) for analysis^23^. As described below, the normal logarithm (ln) of NSAF values was computed to create a more Gaussian/normal distribution, and the Student’s T-Test was utilized to determine the significance of the difference in abundance between experimental and negative control conditions for each potential interacting protein.

### Data analysis

Raw files were searched with no prior peaklist selection by both ProteinMetrics Inc. Byonic (v3.8.13) and ThermoFisher Scientific Proteome Discoverer – SequestHT (2.2.0.338). The proteomic database consisted of all SwissProt annotated human protein sequences (obtained 09-2019 – 20,434 sequences), plus a list of common contaminants (trypsin, keratins, and serum albumins – 179 sequences, available in Supplementary Table 1) and the sequences for eGFP and BirA*. A concatenated database was generated for FDR calculations by including reversed protein sequences for all proteins in the database, creating a final database of 41230 sequences, all of which were searched unbiasedly. Tryptic cleavage was fully specific at Arg and Lys, with two missed cleavages allowed. For both search engines, precursor mass tolerance was 20ppm and fragment mass tolerance was 0.5Da. Carbamidomethylation on Cys was set as a fixed modification, and variable modifications were allowed: Oxidation of Met, HexNAc on Ser/Thr, and Biotin on Lys. Full peptide lists were exported from PMI-Byonic and SequestHT with no score cuts.

Peptide lists from each sample (four fractions each) were then combined using ProValt^24^. Peptide cuts were made based on peptide FDR, set at 2% for the eGFP-BirA* and 1% for TPR-BirA* – with the lower stringency of filtering for eGFP-BirA* samples selected to help decrease the incidence of potential false positives in the final protein set. Protein lists were generated by removing all peptides not identified by both Byonic and Proteome Discoverer (peptides required to have a positive Byonic Score and Sequest Xcorr), and by removing any proteins that were identified by fewer than two peptides. Only proteins identified as the top protein from among isoforms were kept in the final protein list. For initial quantification and putative TPR interacting protein list generation, ln(NSAF) values were calculated for each identified protein and compared between experimental condition and negative control using the Microsoft Excel t.test function. Protein IDs with a pvalue of 0.05 or less and with at least a 3-fold higher NSAF were kept in the final TPR-BirA* interactome.

Further quantification was carried out by reconstructed ion chromatogram (RIC) analysis of peptide intensity performed in Xcalibur Qual Browser (v2.0.3.2). Peptides for analysis were selected based on their appearance in all three replicates in both TPR-BirA* and eGFP-BirA*. Peptide intensity was examined for all gel fractions in which that peptide occurred (For HCF1: fractions 1 and 2; for OGT: fraction 1 [rep 3] and fraction 2 [reps 1,2]; for KNL1: fraction 1). Time ranges for intensity analysis were selected to be the same in TPR-BirA* and eGFP-BirA*. The time range was selected based on the area of overlap between the corresponding peaks, or, in the case of peaks with a slight time offset leading to no overlap, the time range was selected so that the time was evenly split between the two peaks. Peptide intensity was determined by the normalization level (NL) of the monoisotopic peak. Peptide identity was validated by recorded retention time in ProteomeDiscoverer 2.2.

### Western Blots and Antibodies

SDS-PAGE gels (4-15%) were run using the BioRad Mini-PROTEAN gel system. Gels were transferred onto Immobilin-FL PVDF membranes (Sigma) using the BioRad Trans-Blot SD SemiDry Transfer Cell. Membranes were blocked in 1% cold water fish skin gelatin (Sigma), then incubated with primary antibody at these ratios: anti-FLAG (1:2500, Sigma F3165), anti-biotin (1:1000, Jackson 200-002-211), anti-OGT (1:1000, Santa Cruz sc-74546), anti-αtubulin (WB 1:5000, Cell Signaling 3873) Histone H3 (1:1000, Cell Signaling 14269), anti-GAPDH (WB, Cell Signaling 2118). Secondary antibodies were Li-Cor IRDye: 680RD donkey-anti mouse 680 (1:10000), 800CW Goat anti-rabbit (1:20000). Three washes in TBST (0.1% tween) were performed after each antibody incubation. All Western blots were imaged on a LiCor Odyssey Clx system. Densitometric measurements were made using Image Studio Lite v5.2

### Localization studies (Nucleocytoplasmic Fractionation)

Protein localization was determined using UniProt^25^. Nuclear and cytoplasmic fractions from HeLa cells were obtained using subcellular fractionation as previously described^26^ and analyzed via western blot as above.

### Pathway Analysis

Gene ontology analysis was performed using The Gene Ontology Resource (geneontology.org)^27,28^. All GO lists were filtered at pvalue less than 0.01, FDR of less than 0.01, and a minimum of 5-fold enrichment over expected number of proteins found in that category in a random protein dataset. Biological process and molecular function analysis were performed using the GO Ontology Database Released 2019-12-09. ReViGo^29^ was used to generate condensed lists of GO terms and CirGo^30^ to generate plots from the condensed data. PANTHER pathway analysis was performed using PANTHER version 15 released 2020-02-14. Reactome data was also obtained from The Gene Ontology Resource, using Reactome version 65 released 2019-12-22, and parsed at FDR less than 5E-9. Condensed GO term lists and the full reactome pathway list are available in Supplementary Table 5.

Disease association for proteins was identified using the OMIM catalog^31^. Disorders were categorized manually, where “Intellectual Disability” refers to any disorder with the symptom intellectual disability (or several other related terms), “Immunodeficiency” refers to disorders causing immunodeficiency, “Malignancy” refers to any of several cancers, “Congenital, other” refers to congenital disorders not featuring intellectual disability, “Neurological, other” refers to non-congenital neurological disorders, and “Hormone” refers to disorders of the endocrine system.

## Results

### Defining the OGT TPR Interactome in HeLa Cells

To identify OGT TPR interactors, we utilized a fusion protein strategy using promiscuous biotin ligase BirA*. We generated a fusion protein TPR-BirA*, essentially replacing the catalytic domain of OGT with BirA*, and also created a eGFP-BirA* fusion protein to serve as a negative control for nonspecific protein interactions or promiscuous labeling (Supplementary Table 1). Each fusion protein was transiently expressed in HeLa cells and induced with biotin for 24 hours for labeling of proximal proteins (**Fig. 1A**). Note that for transfections, 10x more TPR-BirA* plasmid was used than eGFP-BirA*, due to eGFP-BirA* expressing at a much higher level than TPR-BirA* (**Fig. 1C/D**). Following labeling, we isolated biotinylated proteins with a biotin immunopurification (**Fig. 1E**). It is noteworthy that even though there is significant biotin labeling in both TPR-BirA* and eGFP-BirA*, the band pattern differs significantly indicating a change in the specificity of biotinylation between eGFP-BirA* and TPR-BirA*. A sectioned SDS-PAGE gel was subjected to in-gel digestion and the resulting peptides separated by nanoflow reverse-phase liquid chromatography in-line to a tribrid mass spectrometer for protein identification (see methods). This entire procedure (transfection to LC-MS/MS analyses) was carried out in 3 independent biological replicates for both TPR-BirA* and eGFP-BirA*.

For analysis of the mass spectrometry data, we opted for a multi-algorithm search to increase the stringency of our protein IDs (**Fig. 1B**). Raw mass spectrometry data was searched using both PMI-Byonic and Sequest HT (through ProteomeDiscoverer 2.2) against the human database (Swissprot 09/2019), and only peptides identified by both algorithms were used to generate the final protein set. The negative control (eGFP-BirA*) protein set was searched at a looser peptide FDR (2%) than the TPR protein set (1 %) to further enhance the stringency of the final protein I Ds.

To generate proteins for a final TPR interactors list, we combined uniquely identified proteins in the TPR-BirA* analyses with those that were enriched in the TPR-BirA* protein lists as compared to the eGFP-BirA* (negative control) protein lists. This enrichment was required to be significant according to the Student’s t-test with the cut-off for significance being a p-value of 0.05, and we also required the average NSAF^23^ to be at least 3 times higher in the TPR-BirA* condition compared to the eGFP-BirA* condition. These proteins represent a stringent list of 115 high confidence OGT TPR interactors (**Table 1**, Supplementary Table 4).

**Table 1:**
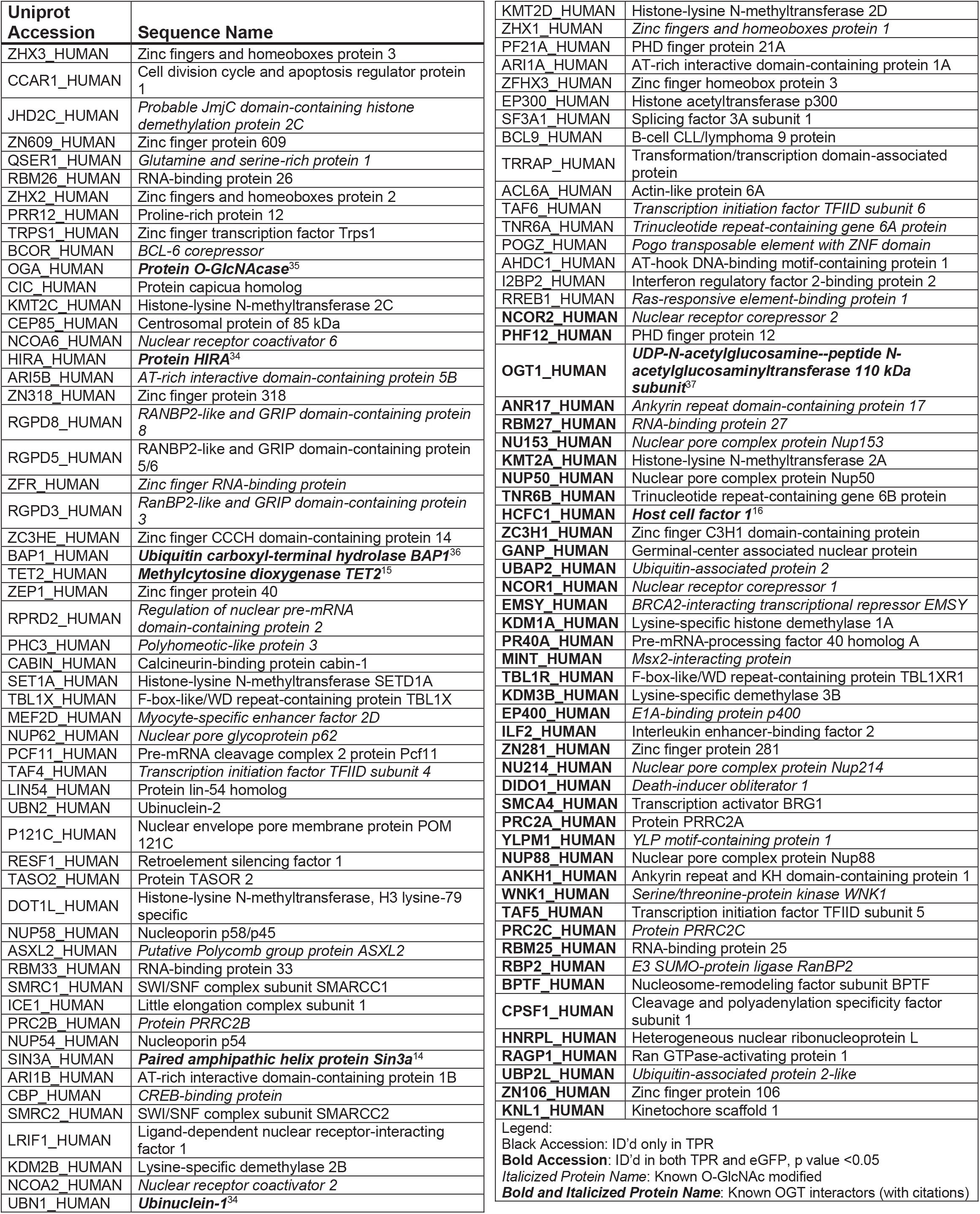
Interactors identified as TPR-BirA* interactors in HeLa cells

46 of the proteins (indicated by an *italicized* protein name in Table 1) had already been identified as O-GlcNAc modified by previous-omic datasets^32,33^. This result suggests that the TPR domain alone can select OGT substrate proteins without the presence of the catalytic domain. In addition, 8 (indicated by an ***italicized and bold*** protein name in Table 1) of the identified proteins had previously been demonstrated to specifically interact with OGT, where “interaction” here is defined as a one or two directional co-immunoprecipitation^14–16,34–37^. Together, these factors lend confidence to the novel protein IDs in the dataset.

It is noteworthy that OGT itself is identified in the screen. Although many peptides attributed to OGT are due to the overexpression of the TPR-BirA* fusion protein, several high-confidence peptides in the catalytic domain of OGT were also identified. This indicates that endogenous OGT, which normally exists as a dimer^11^, did complex with the fusion TPR protein. Several well-studied OGT interactors were also identified, including HCF1^16^, mSin3a^14^, and Tet2^15^. We also identify the O-GlcNAc hydrolase OGA, which OGT is known to regulate both pre- and post-translationally^6,38^.

Many of the interactors identified here are members of protein complexes which may imply that OGT does not directly interact with all of the proteins in the complex. One such complex is the HIRA protein complex, previously shown to interact with OGT^34^. We identified all three members of this complex (HIRA, UBN1, and CABIN). We also identified a novel TPR interaction with the SWI/SNF complex. Six members of the SWI/SNF complex were identified (SMRC2, SMRC1, ACL6A, SMCA4, ARI1A, ARI1B). SWI/SNF proteins, like OGT^39^, function in chromatin remodeling^40^, but OGT has, to our knowledge, never been shown to interact with these proteins. In addition to many protein interactors involved in known OGT functions, we also identified proteins with roles in RNA processing, an area of cellular biology for which limited research exists on the role of OGT and the O-GlcNAc modification. These interactors include proteins with known and putative roles in pre-mRNA splicing (SF3A1, PCF11, PRC2A, PR40A), polyadenylation (CPSF1), and RNA binding (ZN106, TNR6B, RBM33, RBM25, RBM26).

### Validation of Proteins Identified in both TPR-BirA* and eGFP-BirA*

72 of the 115 OGT TPR interactors were only observed in the TPR interactome. Several protein IDs (43 – indicated by a **bold** Uniprot accession in Table 1) were identified in both the TPR-BirA* and eGFP-BirA* samples, but were significantly enriched in TPR-BirA* at the level of average NSAF (Student’s t-test p value < 0.05, fold enrichment of average NSAF >3) (**Fig. 2A**). To further confirm the validity of the inclusion of these proteins in the final dataset, we examined MS1 reconstructed ion chromatograms for peptides identified in both TPR-BirA* and eGFP-BirA*. OGT itself was identified in both, although it is highly enriched in the TPR-BirA*, likely in part due to the overexpression of the TPR-BirA* fusion protein. To confirm that endogenous OGT labeling is enriched in the TPR-BirA* sample, we compared the intensity of a catalytic domain peptide between the TPR-BirA* samples and the eGFP-BirA* samples. The average intensity of this peptide in the TPR-BirA* samples is 9.1 times higher than in the eGFP-BirA* samples, supporting specific interaction of TPR-BirA* with full-length endogenous OGT (**Fig. 3A**).

**Figure 2:**
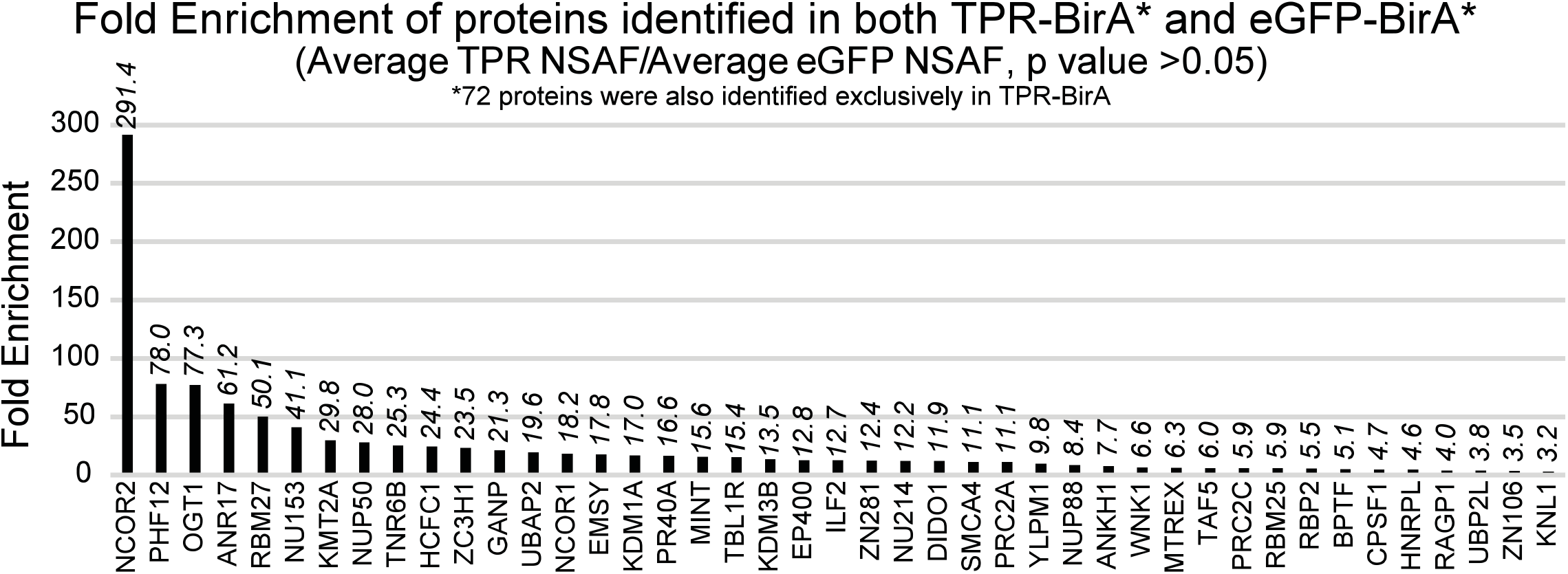
Proteins identified in both TPR-BirA* and eGFP-BirA* are enriched in TPR-BirA*. Fold enrichment values for all proteins identified in both TPR-BirA* and eGFP-BirA*. Fold enrichment values are average NSAF of TPR-BirA* over average NSAF of eGFP-BirA*. Note that 72 proteins were only observed in TPR-BirA*.

**Figure 3:**
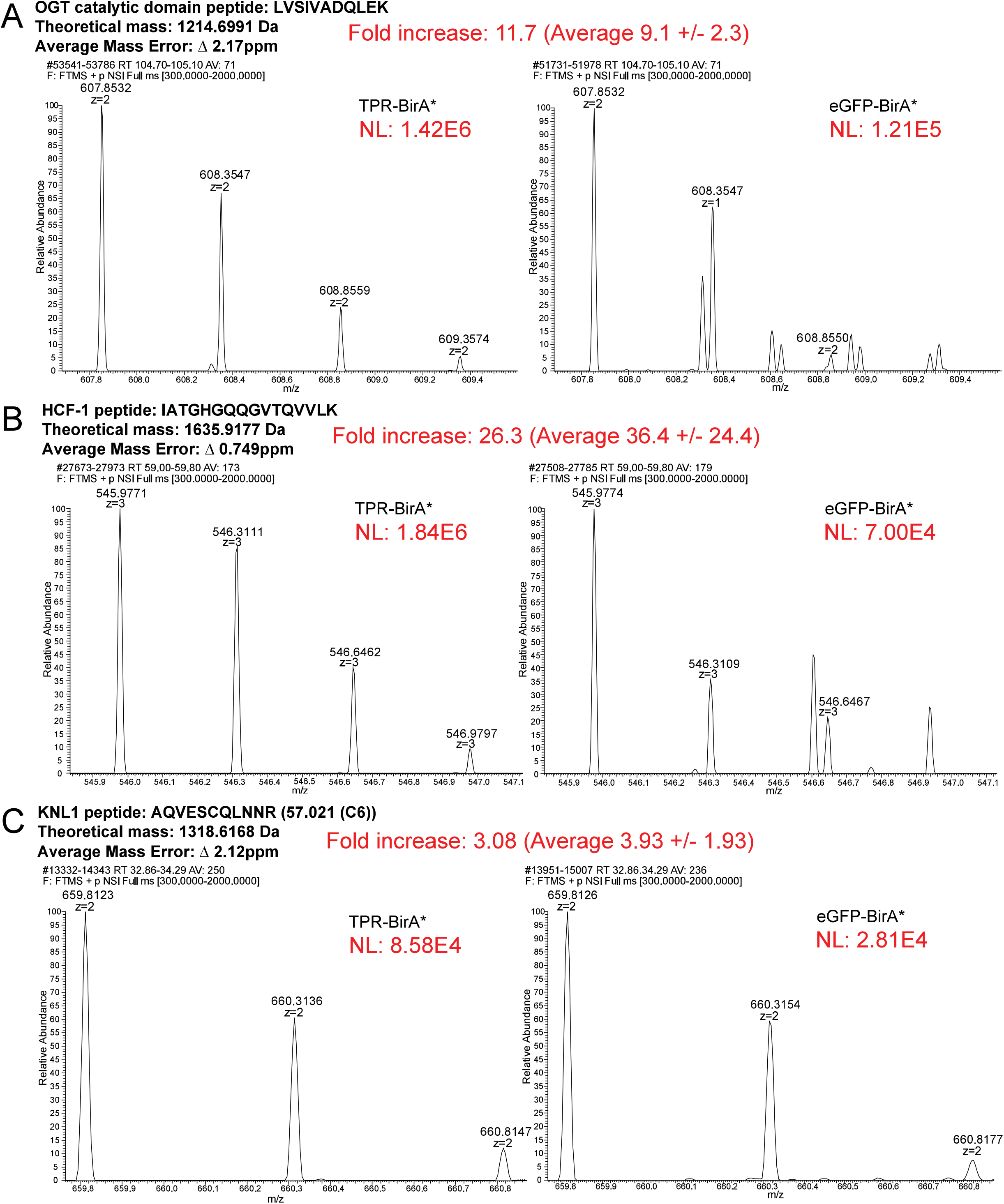
Reconstructed ion chromatograms confirm peptide-level enrichment of TPR-BirA* interactors. **For A-C**, fold increase is for the replicate shown. The average fold intensity for that peptide (averaged across all replicates and fractions in which that peptide appeared) is shown with the standard deviation. Average mass error is the absolute value of the mean across 3 replicates of both TPR-BirA* and eGFP-BirA*. NL=Normalization Level (Base Peak Intensity) ***A***, Reconstructed ion chromatograms demonstrating greater enrichment of a catalytic OGT peptide in TPR-BirA* compared to eGFP-BirA*. ***B***, Reconstructed ion chromatograms demonstrating greater enrichment of an HCF-1 peptide in TPR-BirA* compared to eGFP-BirA*. ***C***, Reconstructed ion chromatograms demonstrating greater enrichment of a KNL1 peptide (novel interactor closest to cut-off for assignment) in TPR-BirA* compared to eGFP-BirA*.

To further confirm the enrichment of relevant OGT interactors in TPR-BirA*, we next examined the intensity of peptides from HCF-1, a well-studied OGT interactor with roles in intellectual disability^16,41^. Average intensity for an HCF-1 peptide are 36.4-fold higher in TPR-BirA* than eGFP-BirA* (**Fig. 3B**). Finally, we confirmed a fold increase in peptide intensity for the protein closest to our fold enrichment cutoff, KNL1 (**Fig. 3C**). The peptide shown from KNL1 has an average intensity in TPR-BirA* that is 3.93 higher than in eGFP-BirA*. Taken together, these results indicate that although some proteins were identified in both the experimental and negative control conditions, their inclusion in the final protein interactor list due to higher enrichment is supported by the raw mass spectrometry data.

### Subcellular Localization of TPR-BirA* Interactors

OGT localizes primarily to the nucleus, but also to the cytoplasm, in the mammalian cell^42^. To confirm that the TPR-BirA* fusion protein also localized to both compartments, we examined the subcellular localization of OGT and our fusion proteins. By subcellular fractionation (**Fig. 4A/B)**, endogenous OGT localizes primarily to the nucleus with some expression in the cytoplasm, as expected^42^. In contrast, the TPR-BirA* fusion protein localizes more highly to the cytoplasm, although it is also present in the nucleus. This result is unsurprising as previous research has shown that overexpressed OGT localizes more highly to the cytoplasm than native OGT^42^. eGFP-BirA* also localizes to both the nucleus and cytoplasm. The subcellular localization profiles of TPR-BirA* and eGFP-BirA* are very similar, supporting the use of eGFP-BirA* as a sufficient negative control for nonspecific labeling by BirA* in both the nuclear and cytosolic compartments. We expected most TPR-BirA* interactors to be primarily nuclear, as most recorded OGT interactors are as well^1^. Analysis of the subcellular localization of identified TPR-BirA* interactors supports this (Visualized in an UpsetR plot **Fig. 4C**, and in a pie chart in **Fig. 4D**). 67 of the 115 identified interactors are exclusively nuclear, with an additional 23 occurring in both the nucleus and the cytoplasm. Several interactors localize specifically to the nuclear pore. Only four proteins exclusively localize to the cytoplasm, all of which are novel OGT interactors (RGPD5, WNK1, TNR6B, and ANKH1). Taken together, this supports the physiologic relevance of the identified TPR interactors and is consistent with prior studies suggestion that OGT interacts primarily with nuclear proteins^1^.

**Figure 4:**
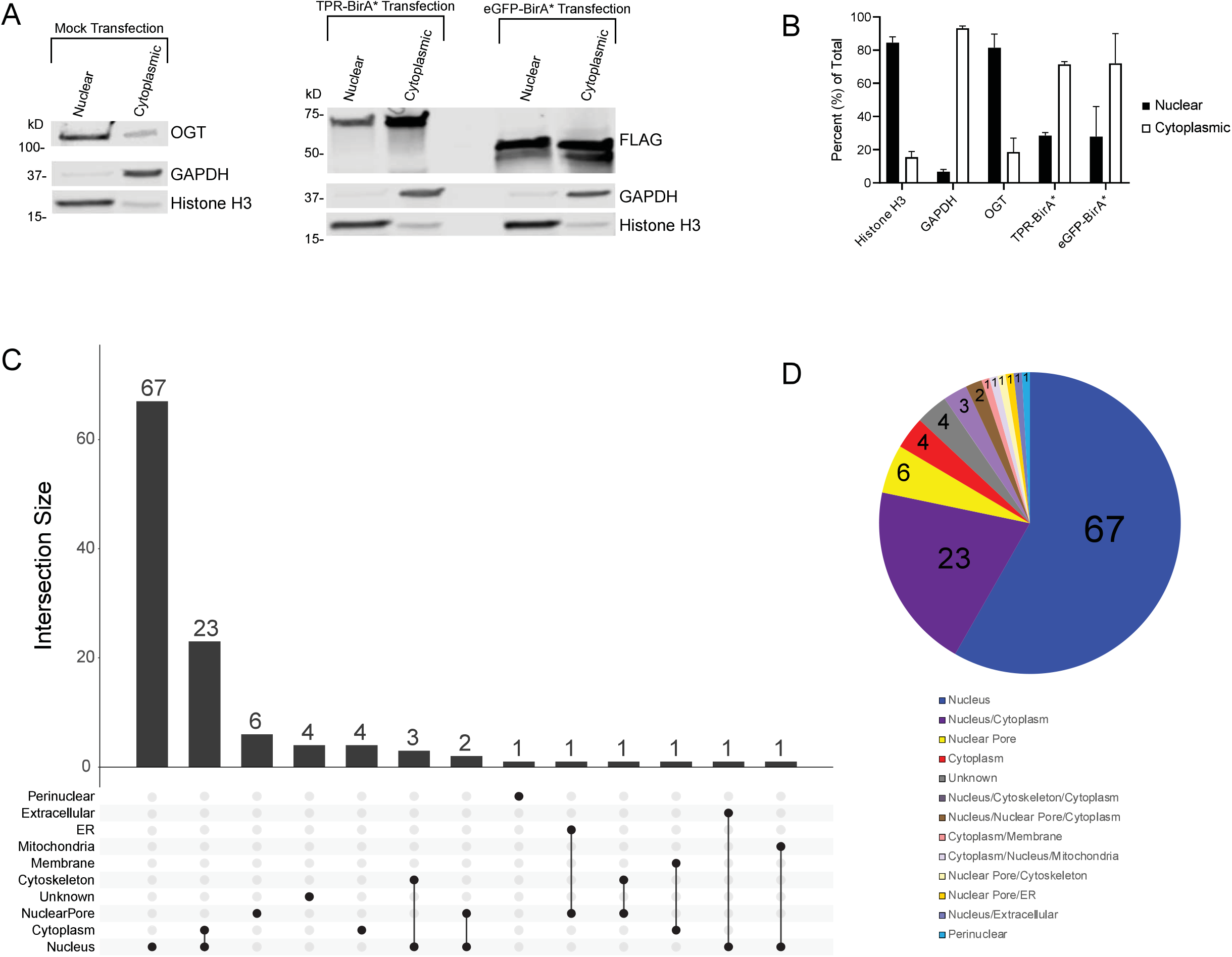
TPR interactors are primarily nuclear localized. ***A***, Subcellular fractionation of HeLa cells demonstrating localization of OGT (anti-OGT F12) and BirA* fusion proteins (anti-FLAG tag). Cytoplasmic marker is GAPDH, nuclear marker is Histone H3. 10ug/lane, representative western blot of three biological replicates ***B***, Ratios of nuclear to cytoplasmic expression of marker proteins (Nuclear: Histone H3, Cytoplasmic: GAPDH) and fusion proteins. Averaged across three biological replicates. ***C***, UpsetR plot showing the subcellular localization of TPR interactors. ***D***, Venn diagram showing the subcellular localization of TPR interactors. Numbers represent the total number of TPR interactors in that category. Localization determined using UniProt.

### Ontology Analyses of the OGT TPR-Interactome

To further understand the enrichment of various processes in our OGT TPR interactome, we performed several different Gene Ontology (GO) analyses using the Gene Ontology Resource (GeneOntology.org) (**Fig. 5**). Performance of a PANTHER Overrepresentation Test for biological processes in our interactome further confirms OGT interactors having frequent roles in transcriptional and chromatin regulation (**Fig. 5A**). General chromatin organization is a strongly enriched category, along with the related peptidyl-lysine modification (indicative of histone modification). Of note is the enrichment of OGT interactors specifically involved in gene silencing. OGT is a Polycomb Group Protein (*sxc* in *Drosophila melanogaster*), which is responsible for the silencing of Hox genes during developmental patterning^43,44^. The interactors identified here further confirm OGT’s tendency toward roles in gene silencing and may reveal further avenues by which OGT regulates gene silencing. Of additional note is the enrichment of interactors involved in the regulation of cellular response to heat, since previous work has demonstrated a role for OGT in coping with cellular heat shock^45^; however, limited work has been published exploring the specific OGT interactions that help it to perform this function. Finally, the enrichment of proteins involved in rhythmic process and circadian rhythm aligns with previous research demonstrating that OGT is involved in circadian rhythm regulation^46^. To confirm these enriched processes, we also performed a PANTHER Overrepresentation Test for molecular function pathways in our OGT TPR interactome (**Fig. 5B**). Many molecular functions corroborate our findings of enriched biological processes, including consistent high enrichment in chromatin and transcriptional regulation. It is interesting to note that RNA Pol II transcription factor binding in particular is an enriched molecular function, since OGT is known to interact with and regulate RNA Pol II-mediated transcription^13,47,48^. Also enriched are processes specifically relating to histone modification, further confirming the enrichment of peptidyl-lysine modification of histones as identified in biological process enrichment and consistent with the O-GlcNAc modification being part of the histone code^49^. Finally, enrichment of nuclear pore components and nuclear localization sequence binding confirms the long-standing role for OGT in nuclear pore structure and/or regulation^50^.

**Figure 5:**
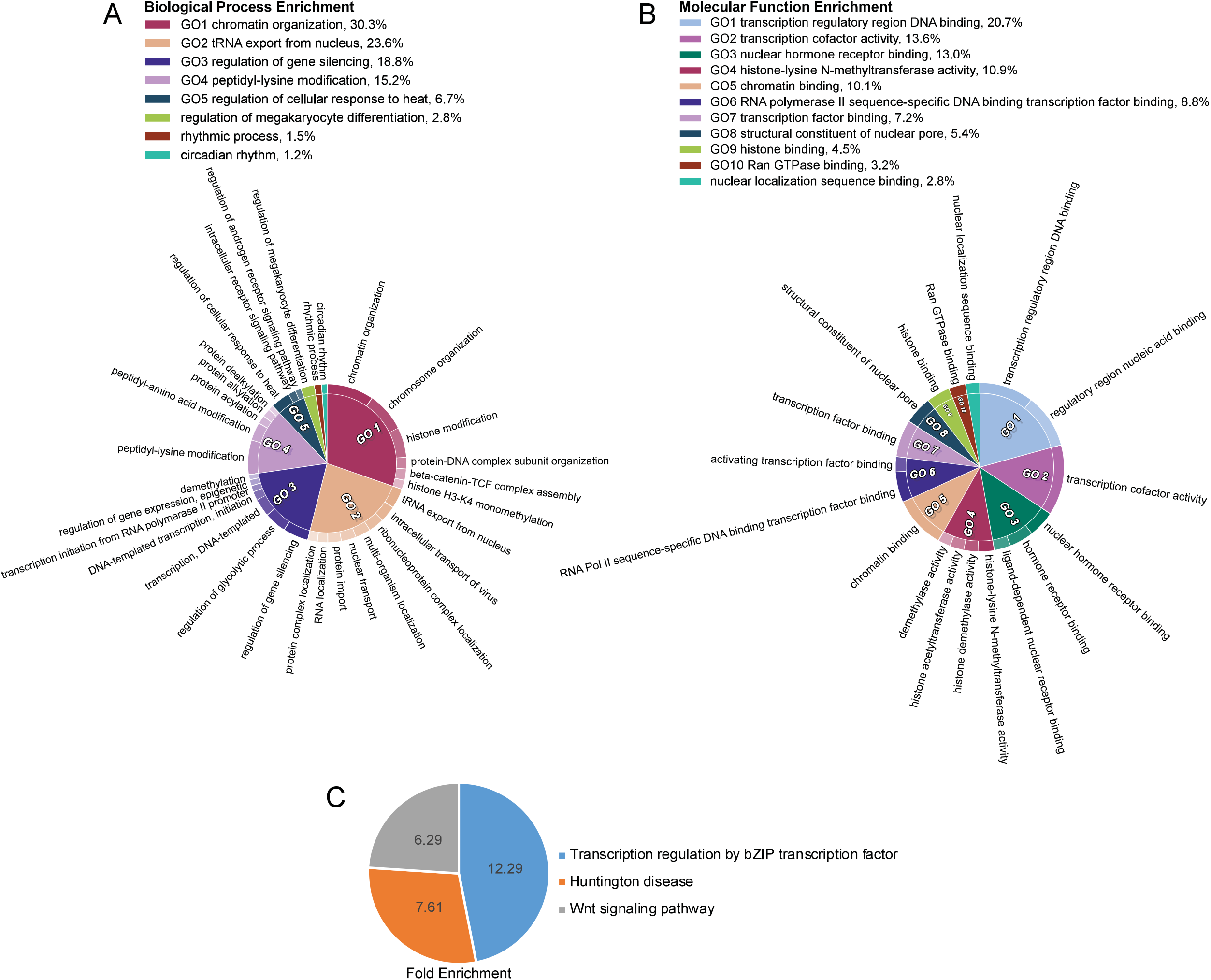
TPR interactors demonstrate enrichment in biological processes and disease states. ***A***, CirGo plot showing enriched biological processes in the TPR protein interactor list ***B***, CirGo plot showing enriched molecular functions in the TPR protein interactor list ***C***, Venn diagram of PANTHER Pathway enrichment of TPR interactors. Numbers are the fold enrichment of the pathway process over expected enrichment.

We further examined enriched Reactome pathways (**Table 2**) among TPR interactors, specifically the mostly highly enriched pathways at an FDR of less than 5E-9. This pathway analysis validates our previous GO analyses that reveal roles for OGT in chromatin regulation, transcriptional regulation, and nuclear pore processes. Reactome pathway enrichment also uniquely reveals several roles for OGT TPR interactors in viral infection, nuclear import, and processing. OGT has been demonstrated to play a role in a limited number of specific viral infections^38,51^ but these enriched pathways point to a potentially broader role for OGT and its interactors more generally in viral infection. Interactors are also enriched in the reactome pathway “regulation of glucokinase by glucokinase regulatory protein”. OGT has already been shown to regulate glucokinase^52^ as well as other proteins involved in glucose metabolism including phosphofructokinase 1^53^. Panther pathway enrichment analysis (**Fig. 5C**) reveals OGT interactor involvement in basic leucine-zipped transcription factor mediated transcriptional regulation, the Wnt signaling pathway, and Huntington disease related processes. The interplay with basic leucine-zipped transcription factors points to another potential avenue for OGT’s regulation of transcription. Furthermore, OGT has already been shown to interface with the Wnt pathway by modulating β-Catenin stability^54^. The interactors identified here involved in this pathway may point to other mechanisms by which OGT modulates Wnt signaling.

**Table 2:**
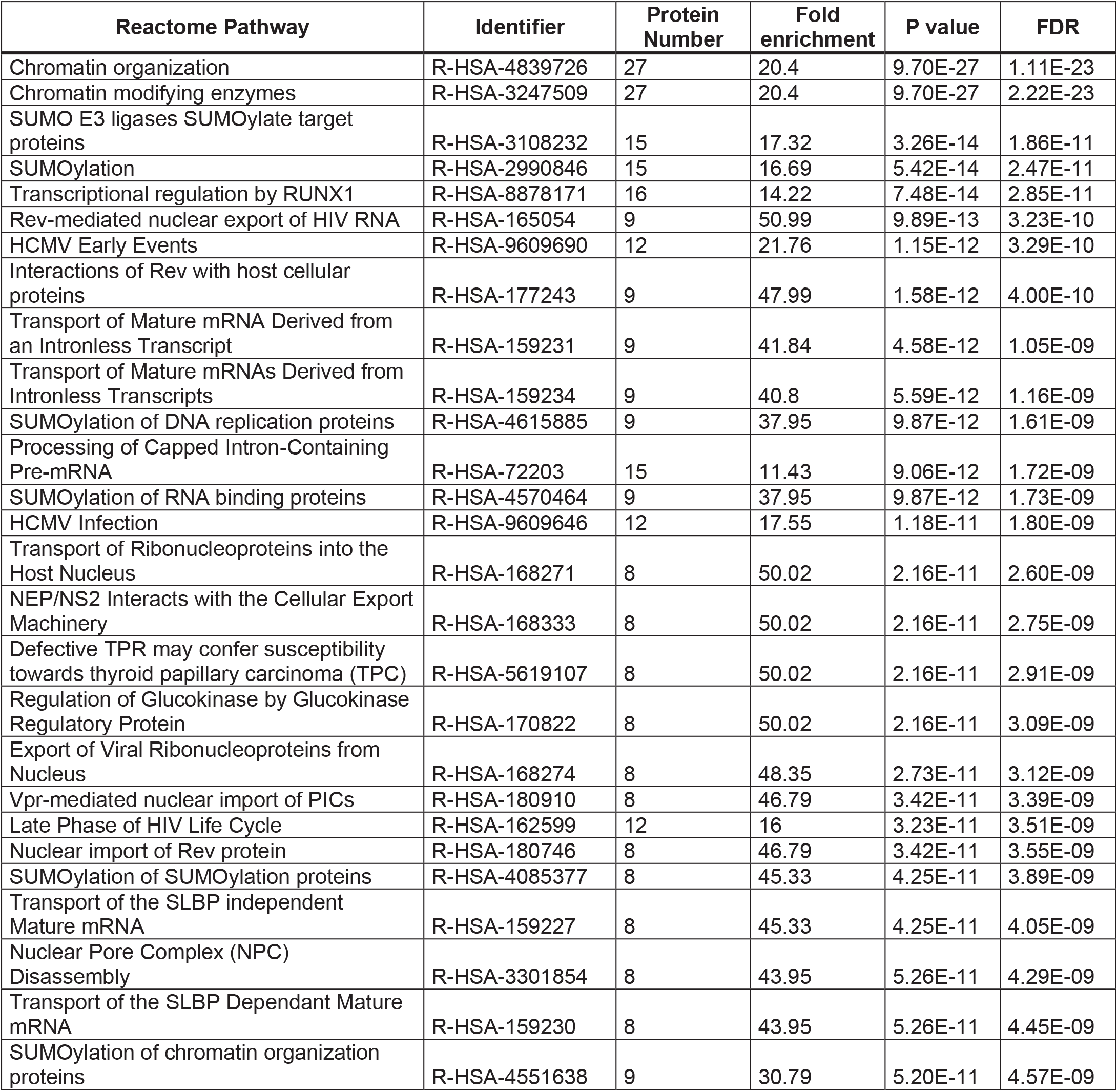
Reactome pathways enriched in TPR interactors list (FDR <5E-9)

| Reactome Pathway | Identifier | Protein Number | Fold enrichment | P value | FDR |
| --- | --- | --- | --- | --- | --- |
| Chromatin organization | R-HSA-4839726 | 27 | 20.4 | 9.70E-27 | 1.11E-23 |
| Chromatin modifying enzymes | R-HSA-3247509 | 27 | 20.4 | 9.70E-27 | 2.22E-23 |
| SUMO E3 ligases SUMOylate target proteins | R-HSA-3108232 | 15 | 17.32 | 3.26E-14 | 1.86E-11 |
| SUMOylation | R-HSA-2990846 | 15 | 16.69 | 5.42E-14 | 2.47E-11 |
| Transcriptional regulation by RUNX1 | R-HSA-8878171 | 16 | 14.22 | 7.48E-14 | 2.85E-11 |
| Rev-mediated nuclear export of HIV RNA | R-HSA-165054 | 9 | 50.99 | 9.89E-13 | 3.23E-10 |
| HCMV Early Events | R-HSA-9609690 | 12 | 21.76 | 1.15E-12 | 3.29E-10 |
| Interactions of Rev with host cellular proteins | R-HSA-177243 | 9 | 47.99 | 1.58E-12 | 4.00E-10 |
| Transport of Mature mRNA Derived from an Intronless Transcript | R-HSA-159231 | 9 | 41.84 | 4.58E-12 | 1.05E-09 |
| Transport of Mature mRNAs Derived from Intronless Transcripts | R-HSA-159234 | 9 | 40.8 | 5.59E-12 | 1.16E-09 |
| SUMOylation of DNA replication proteins | R-HSA-4615885 | 9 | 37.95 | 9.87E-12 | 1.61E-09 |
| Processing of Capped Intron-Containing Pre-mRNA | R-HSA-72203 | 15 | 11.43 | 9.06E-12 | 1.72E-09 |
| SUMOylation of RNA binding proteins | R-HSA-4570464 | 9 | 37.95 | 9.87E-12 | 1.73E-09 |
| HCMV Infection | R-HSA-9609646 | 12 | 17.55 | 1.18E-11 | 1.80E-09 |
| Transport of Ribonucleoproteins into the Host Nucleus | R-HSA-168271 | 8 | 50.02 | 2.16E-11 | 2.60E-09 |
| NEP/NS2 Interacts with the Cellular Export Machinery | R-HSA-168333 | 8 | 50.02 | 2.16E-11 | 2.75E-09 |
| Defective TPR may confer susceptibility towards thyroid papillary carcinoma (TPC) | R-HSA-5619107 | 8 | 50.02 | 2.16E-11 | 2.91E-09 |
| Regulation of Glucokinase by Glucokinase Regulatory Protein | R-HSA-170822 | 8 | 50.02 | 2.16E-11 | 3.09E-09 |
| Export of Viral Ribonucleoproteins from Nucleus | R-HSA-168274 | 8 | 48.35 | 2.73E-11 | 3.12E-09 |
| Vpr-mediated nuclear import of PICs | R-HSA-180910 | 8 | 46.79 | 3.42E-11 | 3.39E-09 |
| Late Phase of HIV Life Cycle | R-HSA-162599 | 12 | 16 | 3.23E-11 | 3.51E-09 |
| Nuclear import of Rev protein | R-HSA-180746 | 8 | 46.79 | 3.42E-11 | 3.55E-09 |
| SUMOylation of SUMOylation proteins | R-HSA-4085377 | 8 | 45.33 | 4.25E-11 | 3.89E-09 |
| Transport of the SLBP independent Mature mRNA | R-HSA-159227 | 8 | 45.33 | 4.25E-11 | 4.05E-09 |
| Nuclear Pore Complex (NPC) Disassembly | R-HSA-3301854 | 8 | 43.95 | 5.26E-11 | 4.29E-09 |
| Transport of the SLBP Dependant Mature mRNA | R-HSA-159230 | 8 | 43.95 | 5.26E-11 | 4.45E-09 |
| SUMOylation of chromatin organization proteins | R-HSA-4551638 | 9 | 30.79 | 5.20E-11 | 4.57E-09 |

### Pathophysiology Analyses of the OGT TPR-Interactome and Orthogonal validation of XLID-related Interactors

The identification of Huntington’s disease (**Fig. 5C**) as an enriched disease process among the TPR interactors prompted us to examine whether identified TPR interactors are involved in other disease processes. Unsurprisingly, as determined using the OMIM catalogue, many TPR interactors are involved in disease processes with which OGT is already associated, including malignancy^55^ and neurological^56^ disorders (**Fig. 6A/B**).

**Figure 6:**
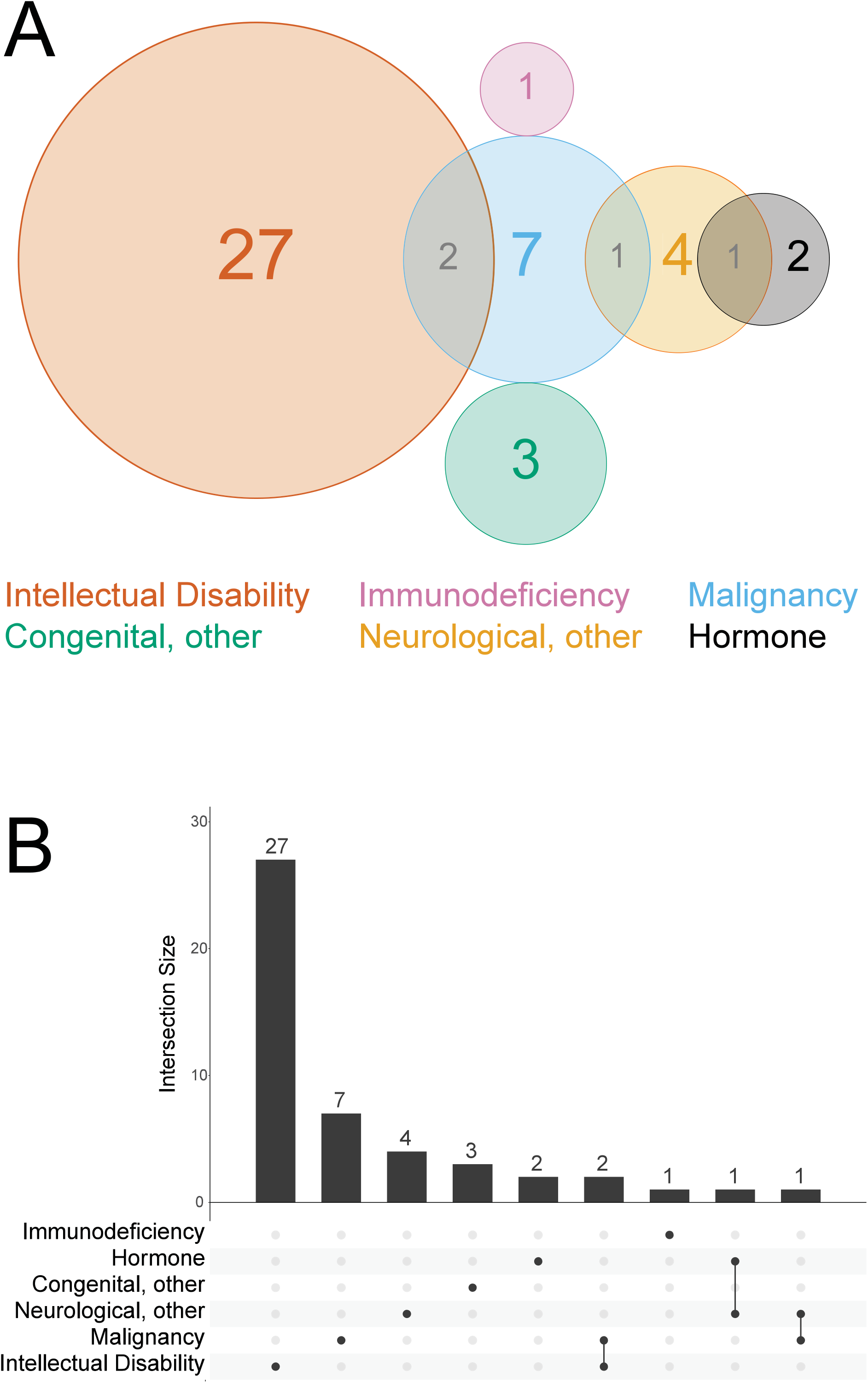
OGT interacts with proteins involved in intellectual disability in HeLa cells. ***A***, Venn diagram showing disease states enriched in the TPR interactors. ***B***, UpsetR plot showing disease states enriched in the TPR interactors. Disease associations determined using the OMIM resource.

Strikingly, of the 44 TPR interactors involved in an OMIM-classified phenotype, 24 are linked to disorders which feature intellectual disability. Three interactors are linked to two different intellectual disability-related disorders each, bringing the total count of intellectual disability disorders associated to OGT interactors to 27 (**Fig 6A/B**). Note that one interactor is also linked to two different malignancy disorders, bringing the total number of disorders associated to TPR interactors to 48. Considering that OGT has recently been linked to X-Linked Intellectual Disability^6–9^, and that the majority of confirmed XLID-causing mutations occur in the TPR domain, these interactors are of high interest in assisting with the elucidation of the mechanism behind variants of OGT being causal for XLID. Furthermore, several of the XLID-associated OGT variants have been demonstrated to be catalytically normal, leading to the hypothesis that the XLID mutations may interrupt protein-protein interactions^6,7,57^. Therefore, these 24 protein interactors are of significant interest in the search for a mechanism underlying the OGT XLID phenotype.

## Discussion

One of the prevailing mysteries in the O-GlcNAc field is how the O-GlcNAc Transferase (OGT) enzyme is able to select from among thousands of possible substrates given that there is only one gene encoding the protein in the mammalian cell. A prevailing hypothesis in the field is that post-translational modification of OGT and protein-protein interactors are responsible for OGT substrate selection. An existing model is that, unlike phosphorylation specificity that evolved by gene duplication followed by divergence driven by evolutionary selective pressures leading to the hundreds of protein Ser/Thr kinases in the mammalian proteome^58^, the O-GlcNAc modification specificity arose from protein-protein associations, perhaps primarily through the TPR domain of the enzyme, evolving to bring substrates into proximity. This mechanism may be similar to RNA PolII that transcribes all protein-coding genes but is exquisitely regulated by protein-protein associations including the basal transcriptional machinery and transcription factors^59^. The role of the TPR domain of OGT in promoting highly specific substrate selection has been demonstrated in part in structural biology studies^11,12,60,61^, but has only been explored for a limited number of specific substrates. Here, we demonstrated that the TPR domain of OGT is capable of interacting with substrate proteins even without the presence of the catalytic domain, and using the BioID technique we have identified 115 high-confidence TPR interactors, representing both known and novel OGT interactors.

OGT is known to localize to the nucleus and the cytoplasm, but usually primarily resides in the nucleus^42^. The fact that most interactors found here are exclusively localized to the nucleus despite the TPR-BirA* fusion protein being localized more dominantly to the cytoplasm is an intriguing observation, suggesting that OGT more strongly and/or frequently interacts with nuclear proteins regardless of localization. Future work is necessary to biochemically confirm each interaction, as well as to determine the degree to which given interactions are transient or stable, under what conditions they occur, and what functional roles they play in the cell.

It is important to note that the identified interactors fall into several different classes. Many of the identified proteins are O-GlcNAc modified, and these may be the effector substrates by which OGT modulates cellular status. Another group, not mutually exclusive to the first, may represent partner proteins; that is, proteins that interact with the TPR domain of OGT to target it to specific substrates or intracellular regions, thus affecting the substrates OGT can access. These proteins may or may not themselves be functionally O-GlcNAc modified. As an example, Tet2 binds to OGT and directs it to histones, but the O-GlcNAc modification on Tet2 has no observed effect on its function^15^. Finally, it is likely that some of the identified interactors do not directly interact with OGT but rather are members of a complex, a subset of which interact directly with OGT. The Swi/Snf complex is an example of this – we have identified 6 of at least 20 possible subunits in our TPR-BirA* interaction list. It is likely that the TPR domain does not directly interact with all 6 identified components and instead interacts with a subset, but additional complex members are labeled due to the diffusion of the reactive biotin intermediate released by the BirA* protein. This would also explain why we fail to identify the full complex; additional members of the complex may be too distant in space to be biotin labeled. The interactors identified here also help to narrow the pool of possible proteins that are directly interacting with OGT, as opposed to a coimmunoprecipitation which would likely pull down the entire stable protein complex. Further work is required to identify direct versus indirect interactors.

In general, the identified interactors confirm OGT’s role as a high-level regulator of cellular function. OGT has previously been characterized as a “rheostat” rather than a switch^1^, and the data here supports this notion. Most of the TPR interactors we identified are “modulators” themselves, e.g. they are not enzymes with a direct effect on a given substrate, but rather affect cellular physiology at a global level by modulating transcription, protein stability, or transport. This gives a perspective of OGT as a modulator of the modulators; that is, OGT regulates cellular function by making many subtle changes in global regulators, adding up to a more significant functional outcome. One such global regulation avenue is chromatin remodeling, which is a previously known function of OGT and a function in which many of our TPR interactors are involved. While our data does not determine how OGT’s interaction with these chromatin remodelers affects their function, many of our identified interactors are involved in lysine modification of histones, pointing to a potential avenue for OGT’s regulation of chromatin remodeling. Indeed, OGT has already been noted to interact with histone modifying enzymes including HDACs^14^. We are unsure why we did not identify any HDACs in our screen – it is possible that they interact with OGT as a part of a protein complex but at a distance outside of the BirA* biotin labeling radius.

Interestingly, we have also identified proteins involved in survival during cellular heat stress. OGT has already been implicated in cellular survival of heat shock^45^. Many of the interactors that fall into this ontology category are nuclear pore proteins. This may indicate that OGT’s role in heat shock is mediated by its modification of nuclear pore proteins, as previously suggested^62^.

We have also identified interactors involved in biological processes in which OGT has yet to be implicated, most noteworthy in RNA processing and transport. Interestingly, OGA, which removes the O-GlcNAc modification, has previously been shown to localize to the nucleolus^63^, indicating the presence and possible role of O-GlcNAc modified proteins in this subcellular structure involved in RNA processing. Future work will be necessary to determine the specific role OGT plays in these processes.

Finally, we have identified many OGT interactors that are involved in disease. OGT and the O-GlcNAc modification are already known to be involved in many disease states including diabetes, cancer, and neurological disorders^1,2^, but this is often only a correlative connection. The TPR interactors we present here may represent avenues for future research uncovering mechanistic proteins underlying OGT’s role in various disease states. Of high interest in the field right now is the mechanism underlying *OGT* mutations leading to X-Linked Intellectual Disability (XLID). One prominent hypothesis that we have previously suggested^6,7,57^ is that mutations in the TPR domain disrupt OGT protein interactions, leading to downstream developmental effects that lead to the XLID phenotype. Here, we have identified 24 OGT TPR interactors directly involved in disorders with intellectual disability. While it is possible that a novel interactor or set of interactors underlies the OGT-XLID mechanism, these interactors represent a strong set of candidate interactors that may contribute to the phenotype. The high number of interactors with connections to intellectual disability may also indicate that there may be a more global interruption in protein-protein interactions caused by XLID variants in OGT. Rather than one specific interactor failing to interact with OGT and leading to XLID, there may be a more subtle reduction in interaction with many proteins, leading to global neurodevelopmental abnormalities.

Our lab is currently undertaking BioID and immunoprecipitation studies to identify any perturbations in the OGT interactome in neural lines harboring XLID-linked OGT variants. The BioID method described here will be a valuable tool to identify potentially tissue-/cell type-specific TPR interactors that fail to interact with XLID-associated OGT variants. In a more directed approach, intellectual disability-related interactors identified here are being screened for protein interaction with XLID-linked OGT variants to determine if they may represent protein interactors underlying the XLID-OGT phenotype. Thus, the work presented here lays a groundwork for additional studies to understand OGT substrate selectivity and the role of OGT and the O-GlcNAc modification in a plethora of biological processes and human pathophysiology including XLID.

## Abbreviations

BirA*: promiscuous Bifunctional ligase/repressor BirA
CID: collision-induced dissociation
IF: immunofluorescence
IP: immunoprecipitation
NSAF: normalized spectral abundance factor
OGT: O-GlcNAc Transferase
OMIM: Online Mendelian Inheritance in Man
TPR: Tetratricopeptide Repeat
WB: Western Blot
XLID: X-Linked Intellectual Disability

## Acknowledgments

We would like to dedicate this paper to the memory of our beloved colleague Dr. Brent Weatherly, who developed the proteomic analysis workflows used in this manuscript. We thank Dr. Kelley Moremen for technical advice and plasmid constructs. This work was supported by a grant from the W.M. Keck foundation (L.W. Co-PI); an NICHD National Institutes of Health (NIH) grant R21HD097652 (L.W); and an NICHD Grant F30 HD098828 (H.S.). The content is solely the responsibility of the authors and does not necessarily represent the official views of the National Institutes of Health.

## Data Availability

The final TPR interactors list along with statistical analyses is available in supplementary table 2. All peptide matches (supplementary table 3) and protein IDs from each replicate (supplementary table 4) along with GO term lists (supplementary table 5) are attached as supplementary data. Raw mass spectrometry data (24. raw files) is deposited in the MassIVE database (https://massive.ucsd.edu/ProteoSAFe/static/massive.jsp, Dataset ID: MSV000085626).

## Author Contributions

H.S. and L.W. conceived and coordinated the study and wrote the manuscript. H.S. performed all experiments and data searches. J.P. performed calculations and statistical analyses on final protein lists and assisted in revision of the manuscript.

